# BulkRNABert: Cancer prognosis from bulk RNA-seq based language models

**DOI:** 10.1101/2024.06.18.599483

**Authors:** Maxence Gélard, Guillaume Richard, Thomas Pierrot, Paul-Henry Cournède

## Abstract

RNA sequencing (RNA-seq) has become a key technology in precision medicine, especially for cancer prognosis. However, the high dimensionality of such data may restrict classic statistical methods, thus raising the need to learn dense representations from them. Transformers models have exhibited capacities in providing representations for long sequences and thus are well suited for transcriptomics data. In this paper, we develop a pre-trained transformer-based language model through self-supervised learning using bulk RNA-seq from both non-cancer and cancer tissues, following BERT’s masking method. By probing learned embeddings from the model or using parameter-efficient fine-tuning, we then build downstream models for cancer-type classification and survival-time prediction. Leveraging the TCGA dataset, we demonstrate the performance of our method, *BulkRNABert*, on both tasks, with significant improvement compared to state-of-the-art methods in the pan-cancer setting for classification and survival analysis. We also show the transfer-learning capabilities of the model in the survival analysis setting on unseen cohorts.

**Data and Code Availability:** In this paper, we leverage the Cancer Genome Atlas (TCGA, https://portal.gdc.cancer.gov/), which includes bulk RNA-seq samples as well as clinical targets for each patient (cancer type, survival time). For pre-training experiments, this dataset is completed with non-cancerous bulk RNA-seq samples from GTEx (Carithers and Moore, 2015) and ENCODE (de Souza, 2012). Code available at https://github.com/instadeepai/multiomics-open-research

**Institutional Review Board (IRB):** This research does not require IRB approval.

## 1. Introduction

New high-throughput technologies have revolutionized molecular research for personalized medicine by providing extensive genomic and transcriptomic data. These have proved to be essential in oncology to help understand the molecular foundations of the disease and to inform on cancer diagnosis or prognosis at the patient level (Stark et al., 2019; Dai and Shen, 2022), notably for cancer type classification or survival analysis. The Cancer Genome Atlas (TCGA) is a publicly available dataset that gathers multi-omics data such as genome sequencing, RNA sequencing, DNA methylation, or whole slide images from thousands of tumor samples. This dataset, divided into 33 cohorts, also provides clinical information such as survival time, making it a widely used dataset for the benchmark of survival analysis and cancer-type classification methods. Survival analysis, or time-to-event prediction, aims at predicting the time from diagnosis to patient death from the disease using censored data. The classic approach for this task has long been the semi-parametric Cox proportional hazards (CPH) model (Cox, 1972). More recently, deeplearning methods such as *Cox-nnet* (Ching et al., 2018) or *DeepSurv* (Katzman et al., 2018) estimate the log-risk function of the CPH model using the Cox partial likelihood as the loss function. Compared to CPH, these models have the advantage of capturing non-linear patterns and interactions between genes underlying the complexity of RNA-seq data.

However, these methods face the *curse of dimensionality* that comes with the high number of genes considered when using RNA-seq data and the small number of available samples. Though some *a priori* feature selection from domain knowledge (such as highly variable genes selection in the case of RNA-seq data) can be carried out, one tends to first learn representations of such high-dimensional data before applying them to any downstream task. Such representations are often obtained through statistical methods such as Principal Component Analysis (Jolliffe, 2002) or Non-negative Matrix Factorization (NMF) (Lee and Seung, 2000) to take advantage of the positivity of gene expressions.

Nevertheless, deep learning-based methods have progressively emerged as state-of-the-art methods for representation learning and have already proven their capacity when applied to omics data. In Gross et al. (2024), a robust evaluation of deep learning-based methods for survival analysis and gene essentiality prediction is presented, with methods mainly based on auto-encoders or graph neural networks. Benkirane et al. (2023) presents a model called *CustOmics* based on Variational Auto-Encoders (Kingma and Welling, 2013) that combines multiple sources of omics data (RNA-seq, Copy Variation Number, and Methylation) for cancer type prediction and survival analysis. They namely show in an ablation study that RNA-seq is the most informative modality for predicting the phenotype of interest, thus motivating our choice of building a model exclusively based on RNA-seq. Vale-Silva and Rohr (2021) also consider a multi-modal model, including especially RNA-seq and whole slide images for survival analysis. Finally, Wu et al. (2024) leverages multi-task learning using the tabular form of RNA-seq data and obtains good performance in survival analysis with a pre-selection of a small subset of genes. Motivated by their wide success in Natural Language Processing, Transformers (Vaswani et al., 2017) models have been used in a variety of domains (image (Dosovitskiy et al., 2020), audio, etc.) by further empowering self-supervision to build foundation models. Thanks to the large amount of available unannotated data, these models have already had an impact in the computational biology field, especially in the analysis of DNA sequences (Dalla-Torre et al., 2023; Ji et al., 2021). More recently, transformer-based models have been used for transcriptomics data to learn representations of single-cell RNA-seq data, with models like *scBERT* (Yang et al., 2022) and *scGPT* (Cui et al., 2024) with application to cell-type annotation for example, thus suggesting applying the same techniques on bulk RNA-seq data. Indeed, even though single-cell or spatially-resolved transcriptomics are getting more traction in medical research as they provide fine-grained analysis, the cost of such sequencing technologies prevents them from being used in large-scale clinical trials or clinical routine, contrary to bulk RNA-seq data, which is now largely accessible and more suited to macro-analysis (cancer type classification and survival analysis).

In this paper, we present a transformer-based encoder-only language model, *BulkRNABert*, pretrained on bulk RNA-seq data through self-supervision using masked language modeling from BERT’s method (Devlin et al., 2018). We will consider language models trained on cancer data from TCGA, but also models trained on non-cancer datasets, GTEx (Carithers and Moore, 2015) and ENCODE (de Souza, 2012)), both to increase the number of bulk RNA-seq samples available, but also to show that one can get good performance in cancer downstream tasks from non-cancer pre-trained models. The learned RNA-seq embeddings are then used to train separate models for the two downstream tasks that we will consider: cancer-type classification and survival analysis. For survival prediction, we will consider both the per-cohort and the pancancer setting, the latter allowing us to consider more samples, thus stabilizing models’ training and also showing transfer learning to unseen cohorts. Moreover, for the classification problem, we will show that fine-tuning the language model through a parameter-efficient fine-tuning method (PEFT) (Xu et al., 2023) further increases our performance. Our method constitutes the first language model pre-trained on bulk RNA-seq data and tailored for cancer type classification and survival analysis, achieving state-of-the-art performance in the pan-cancer setting.

## 2. Methods

### 2.1. Language model pre-training

Language Models (LM) aim at approximating probability distributions over sequences of tokens. In the context of RNA-seq data, each sequence is composed of real values corresponding to gene expression (in *Transcript Per Million*, or *TPM*, units). The first step is thus to transform each gene expression sequence into a sequence of tokens; this is achieved by binning the TPM values on a linear scale. Thus, the token id associated with a given RNA-seq value is its corresponding bin id, so after tokenization one RNA-seq sample is a vector **X ∈** ⟦ 0 ; *B −* 1 ⟧ ^*N*^, with *N* the number of considered genes and *B* the number of expression bins. Our architecture follows an encoder-only transformer trained using Masked Language Modeling, following a classic *BERT* methodology. Compared to the classic transformer architecture, as gene expression is permutation invariant, positional encoding is here replaced by a gene embedding added to the gene expression embedding. The gene embedding matrix, of size *N × n*_*gene embedding*_, given *n*_*gene embedding*_ the dimension of gene embedding, is initialized with *Gene2Vec* method (Zou et al., 2019) as suggested in Yang et al. (2022). To avoid being restricted to the embedding dimension chosen by *Gene2Vec*, the obtained gene embedding is passed through an additional linear layer to reach effective embedding size (*n*_*embedding*_). The final embedding is then processed through a stack of self-attention layers before being transformed by a language model head to get a probability distribution over the RNA-seq bins for each token. We follow typical parameters for Masked Language Modeling: for a given sequence, 15% of tokens are corrupted and used to train the model. Out of those 15%, 80% is replaced by a special mask token [MASK], 10% of the 15% are replaced by random tokens, and the remaining 10% tokens are kept unchanged, but still counted in the masked language modeling loss. The following loss is then optimized:

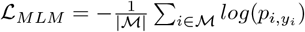

with ℳ *⊂* ⟦ 0 ; *N −* 1 ⟧ is the set of masked token indices, *y*_*i*_ ∈ ⟦ 0 ; *B* −1⟧ is the target gene expression bins and *p* ∈ ℝ^*N ×B*^ is the probability distribution output by the language model.

### 2.2. Cancer type classification

We evaluate our pre-training strategy on several downstream tasks, the first one being a classification task over cancer type. Indeed, cancer classification is vital for accurate diagnosis and therapy, yet classifications based on morphological or histological characteristics are limited, and gene-expression-based methods prove to provide accurate results (Mohammed et al., 2021; García-Díaz et al., 2020). The *TCGA* dataset is divided into 33 cohorts, allowing classification using RNA-seq data with cohorts as labels. The output of the last self-attention layer is extracted and averaged across the sequence dimension and is used as the embedding for a given RNA-seq sample. Therefore, from a raw RNA-seq sample (a vector ℝ^*N*^), one obtains a representation in 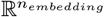. This embedding is then fed to a Multi-Layer Perceptron (MLP), with dropout (Srivastava et al., 2014) and layer normalization (Ba et al., 2016) to get the cancer type prediction.

### 2.3. Survival analysis

The second task is survival time prediction, *i*.*e*. we are interested in building a model that estimates the expected time until the occurrence of an event, in our case, the time until the death of a cancer patient from the time of diagnosis. Thus, we aim to develop a model to predict survival time 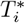 for a given individual *i*. One major issue in survival problems is the concept of right-censoring: the period of observation time 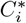 might be less than 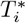 (end of the study, lost contact with the patient). One thus defined 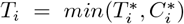 as the target survival time, as well as 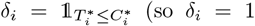 if event occurred (death), otherwise *δ*_*i*_ = 0), thus constituting a dataset 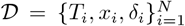, with *x*_*i*_ covariates (RNA-seq mean embeddings in our study). The most popular survival method to solve such a problem is the semi-parametric model called Cox proportional hazard model (Cox, 1972) in which one aims at modeling the hazard function *λ*, defined as the instantaneous event rate at time *t* given survival until time 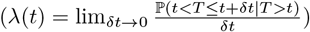 as 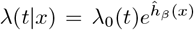, with *β* a vector of parameters (so in our case *x*, 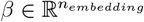), *λ*_0_(*t*) the baseline hazard, and in the Cox model, 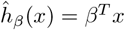. To estimate the vector of parameters, maximum likelihood estimation is used, which is done in the Cox model by maximizing the following partial likelihood:

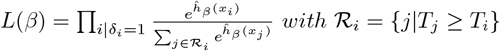

As done in other works, like *DeepSurv* (Katzman et al., 2018) or *Cox-nnet* (Ching et al., 2018), we can replace 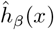 by a neural network (same architecture as for the previous classification head) output and optimize negative partial log-likelihood:

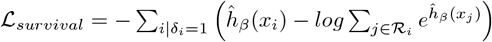

The stability of this training method is dependent on the number of positive examples, and alternatives like *L*_2_-like losses (Jeong and Jia, 2022) could also be explored. This study is left for future work.

### 2.4. *IA*^3^ fine-tuning

For the classification setting, we furthermore employ the *IA*^3^ technique (Liu et al., 2022), a parameter-efficient fine-tuning method that adds new learnable weights to the transformer layers while keeping frozen the pre-trained ones. To this end, three learned vectors 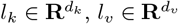, and 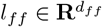 are adjoined into the attention mechanism:

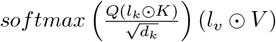

and in the position-wise feed-forward networks through (*l*_*ff*_ *γ*(*W*_1_*x*))*W*_2_, where *γ* is the feed-forward network non-linearity, and ⊙ represents the elementwise multiplication. Compared to a full fine-tuning of the language model and the classification head, this method only adds *L*(*d*_*k*_+*d*_*v*_+*d*_*ff*_) parameters (with *L* the number of self-attention layers), which represents less than 0.07% of the total number of model parameters, thus speeding up the fine-tuning. The same classification head is still used. Complete architecture, from pre-training to task-specific fine-tuning, is presented in Figure 1.

**Figure 1:**
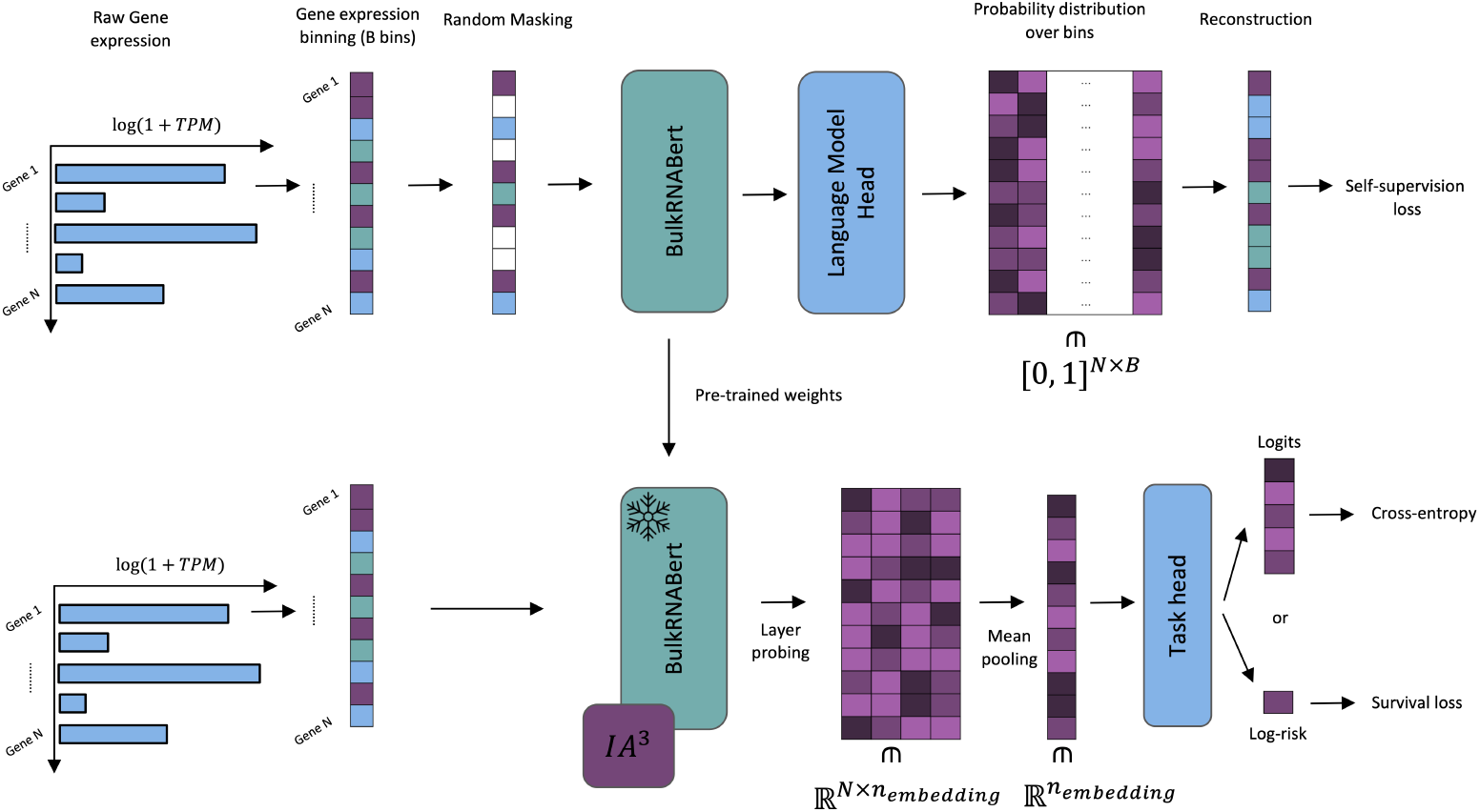
*BulkRNABert* pipeline. The 1st phase consists in pre-training the language model through masked language modeling using binned gene expressions. The 2nd phase fine-tunes a task-specific head using either cross-entropy for the classification task or a Cox-based loss for the survival task. *IA*^3^ rescaling is further added for the classification task.

## 3. Experimental Results

### 3.1. Language Model pre-training

#### Datasets

Three bulk RNA-seq datasets were considered to pre-train *BulkRNABert* : 2 non-cancer datasets, GTEx and ENCODE, representing 20,406 samples, and one cancer dataset, TCGA (all cohorts are considered here), representing 11,274 samples. Among all the genes available in these 3 datasets, we keep all the genes that are common in all 3 datasets, representing a total of *N* = 19,042 genes. We use the transcript per millions (TPM) for the RNA-seq samples unit and apply *x* ↦ *log*_10_(1 + *x*) transformation followed by a max-normalization. The same transformations are applied to RNA-seq samples used in the downstream tasks in the following sections. Using these datasets, we were able to train two language models: *BulkRNABert(GTEx ENCODE)* - pre-trained on GTEx and ENCODE only; *BulkRNABert(TCGA)* - pre-trained on TCGA only. For ablations studies, we additionally pretrained a third model - *BulkRNABert(GTEx ENCODE TCGA)* - pre-trained on the concatenation of GTEx, ENCODE, and TCGA. For the masked language modeling step, each dataset is split into a train and test set, keeping 5% of the dataset for the test set. The masked language modeling loss is optimized using *AdamW* (Loshchilov and Hutter, 2017) optimizer with gradient accumulation to reach an effective batch batch of 3 × 10^6^ tokens, split onto 4 TPUs v4-8.

#### Pre-training

We trained the 3 models using the same set of hyperparameters. Following the presented tokenization method, RNA-seq samples are binned using *B* = 64 bins. Each model is composed of 4 transformer blocks with 8 attention heads each and uses an embedding dimension of 256 (so *n*_*embedding*_ = 256), for a total of 6 million parameters (so less than for example *CustOmics* (Benkirane et al. (2023)) that uses 50 million parameters). Models are trained on a total of 12*B* tokens. Unlike methods like Yang et al. (2022), we decided not to consider only non-zero expressions in masked language modeling but all of them. In Appendix A (Figure 6), we provide a correlation plot that compares the true and the predicted tokens’ distribution to assess that the model is not learning a trivial distribution (*e*.*g* full of zeros). Models are evaluated using the reconstruction accuracy. We provided in Appendix A (Figure 7) the learning curves (in accuracy) of the 3 models: we observe that the reconstruction accuracy is higher when considering the non-cancer datasets GTEx and ENCODE than for TCGA, which can be qualitatively explained by the higher variability of gene expressions in cancerous tissues. Overall, all models exhibit reconstruction accuracies between 0.5 and 0.6 which is a significantly great performance as we are using *B* = 64 bins, *i*.*e*. 64 possibilities for the labels (and 1*/B ≈* 0.016).

### 3.2. Cancer-type classification

We then first assess the quality of the learned representations through cancer-type classification, using TCGA cohorts as class labels. We consider two settings for this classification problem:

- 5 cohorts classification using BRCA, BLCA, GBMLGG, LUAD, and UCEC cohorts (GBMLGG is the fusion of two brain cancer cohorts, GBM and LGG. These are the cohorts considered in Benkirane et al. (2023) for survival analysis, so as a first step, we consider this simpler classification problem using these cohorts), see Appendix B (Figure 8) for label distribution.
- Pan-cancer classification using the 33 cohorts, see Appendix B (Figure 9) for label distribution.

#### Experimental setup

Classification models consist of a MLP with two hidden layers - respectively with 256 and 128 units, using *SELU* (Klambauer et al., 2017) as activation function on top of *BulkRNABert*, followed by a classification head according to the number of classes. We also tried to fit a Support Vector Machine (SVM) (Cortes and Vapnik, 1995) classifier instead of an MLP. We use cross-entropy loss, optimized using Adam (Kingma and Ba, 2014) optimizer. The 5 cohorts classification problem does not involve a significant dataset imbalance, we thus report the weighted *F*_1_ score as an evaluation metric. For the pan-cancer classification, we report both macro *F*_1_ and weighted *F*_1_ scores to avoid any bias due to class imbalance. For each task, the dataset is split into 80% train and 20% test. This is done with 5 different seeds, and the reported score corresponds to the mean score across these 5 seeds. The splits are stratified to ensure equal representation of each class in the train and test splits. Multiple RNA-seq samples might be available for a given patient; thus, we ensure that samples from the same patient are not mixed in the train and test sets.

#### 5 cohorts classification

We compare our method to *CustOmics* (Benkirane et al., 2023) and to two dimension reduction techniques (PCA and NMF) applied to raw RNA-seq and fed to an SVM. We present our results in Figure 2 and exhibit that our method, *BulkRNABert* pre-trained on TCGA and fine-tuned using *IA*^3^ provides the best performance.

**Figure 2:**
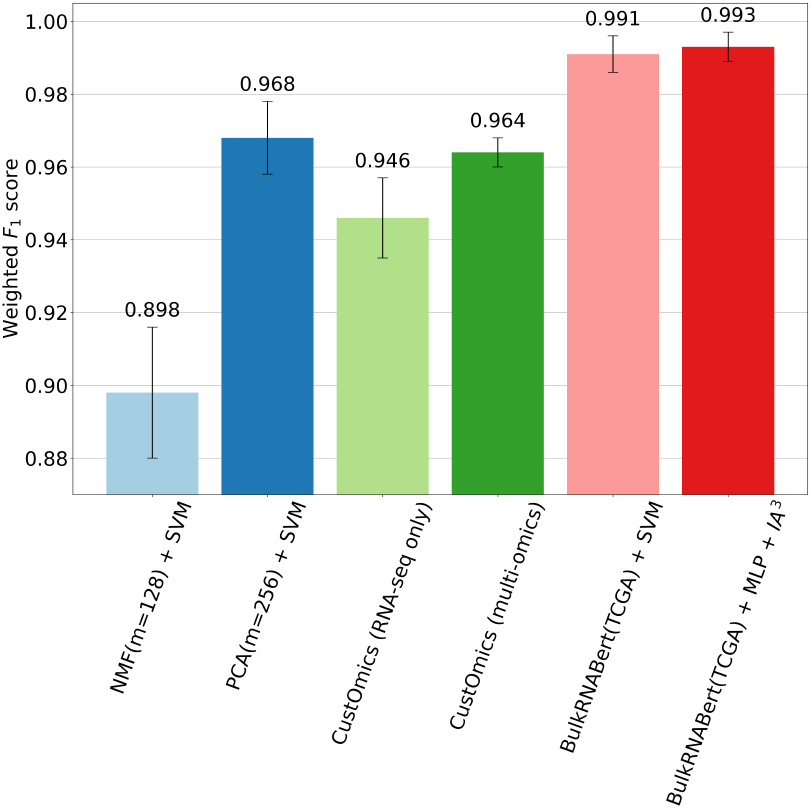
5 cohorts classification results. “*BulkRN-ABert(TCGA)* + MLP/SVM”: only MLP/SVM is trained, while freezing the language model weights. “+ *IA*^3^” indicates that the language models’ weights are fine-tuned with *IA*^3^ method. *m* for PCA and NMF indicates the number of components.

#### Pan-cancer classification

Results are presented in Table 1. Again, we demonstrate that pretraining the language model on TCGA and finetuning it using *IA*^3^ provides state-of-the-art results for cancer-type classification. Further improvements are shown (weighted *F*_1_ of 0.97) by *CustOmics* when adding Copy Number Variation and Methylation. A confusion matrix for our best model is presented in Appendix B (Figure 10), revealing that the remaining errors are mostly due to similar cancers (typically *READ* and *COAD*, respectively linked to rectum and colon).

**Table 1:** Pan-cancer classification. *NA* indicates that the score is not provided by the author.

| Methods | test macro-F1 | test weighted-F1 |
| --- | --- | --- |
| *CustOmics (RNA-seq only)* Benkirane et al. (2023) | *NA* | $0.930 \pm 0.010$ |
| NMF(m=128) + SVM | $0.749 \pm 0.006$ | $0.800 \pm 0.009$ |
| PCA(m=256) + SVM | $0.849 \pm 0.010$ | $0.877 \pm 0.006$ |
| *BulkRNABert(TCGA)* + SVM | $0.902 \pm 0.007$ | $0.933 \pm 0.002$ |
| *BulkRNABert(TCGA)* + MLP + $IA^3$ | **$0.918 \pm 0.006$** | **$0.942 \pm 0.004$** |

For both classification tasks, more experimental results are presented in Appendix B (Tables 5 and 6), showing that pre-training on non-cancer datasets (GTEx and ENCODE) already provides better results than *CustOmics*. Also, we notice that mixing cancerous and non-cancerous tissues in the pretraining does not bring any additional performance, which may root from the same number of parameters kept for the language model, hence a dilution of the data.

### 3.3. Influence of pre-training

To assess the relevance of pre-training compared to directly fitting a randomly initialized model, we compared the following methods on the pan-cancer classification problem:

- **Pre-trained**: language models pre-trained either on TCGA (*Pre-trained (TCGA)*) or GTEx ENCODE (*Pre-trained (GTEx ENCODE)*) with a classification head and *IA*^3^ rescaling while keeping other LM’s weights frozen.
- **No pre-training + full fine-tuning**: same architecture as above but with randomly initialized weights, no *IA*^3^ rescaling and full fine-tuning. We adapted some hyperparameters (weight decay and dropout) to circumvent a performance drop.
- **Pre-trained + full fine-tuning**: language models pre-trained on TCGA with all weights that are fine-tuned.

Results in Figure 3 show that the pre-trained models provide the best performance compared to the randomly initialized and full-fine-tuned model, even when considering a model pre-trained on non-cancerous gene expressions (*Pre-trained(GTEx ENCODE)*). Also, pre-trained models reach their final performance only after a few training steps. We provide in Appendix B (Table 4) the quantitative values for the test macro-F1 of the different models. Also, fully fine-tuning from a pre-trained checkpoint show similar results than using only *IA*^3^ rescaling, hence our choice to go for *IA*^3^ rescaling for memory efficiency.

**Figure 3:**
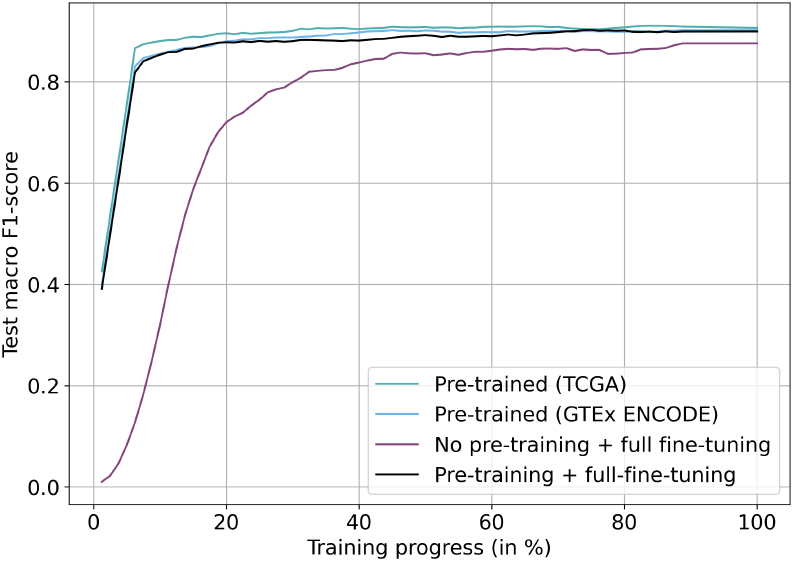
Influence of pre-training on the pan-cancer classification task. Comparison of test macro-F1 score for different pre-training strategies.

### 3.4. *BulkRNABert* embeddings maintain similar cancer types

As a first work on *BulkRNABert* ‘s interpretability, for each cohort, we averaged the *BulkRNABert* embeddings (computed after fine-tuning on the classification task) of every sample belonging to that cohort. Then we computed the cosine similarity between every cohort’s mean embedding and performed a hierarchical clustering presented in Figure 4. One notices that similar cancer types (like GBM/LGG or READ/COAD) have large cosine similarities.

**Figure 4:**
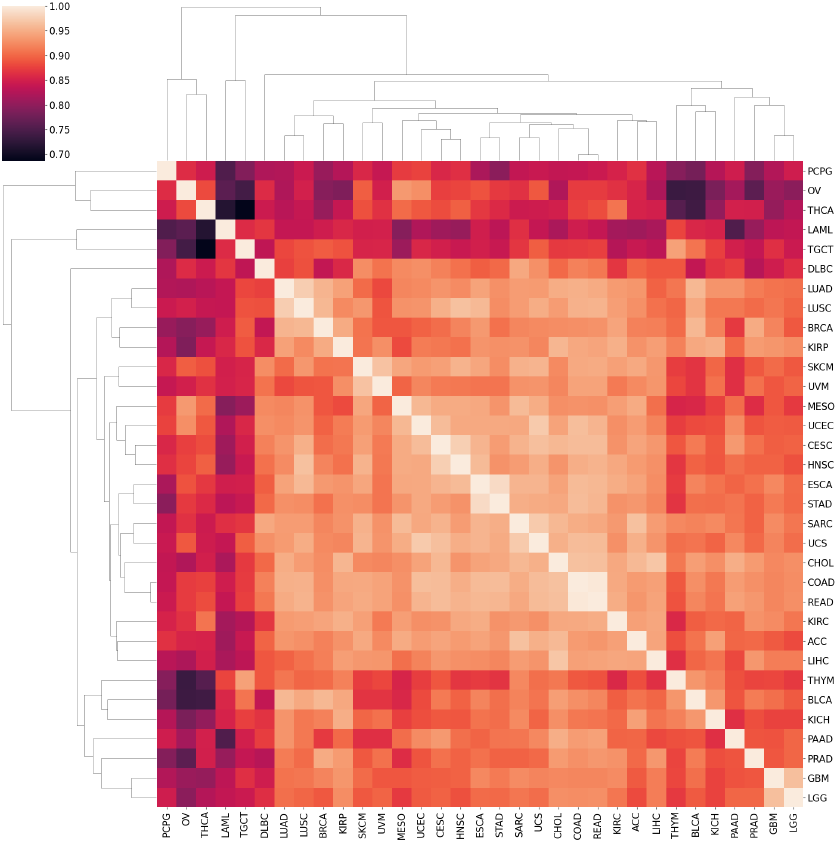
Clustering of cohort’s mean embedding based on cosine similarity. Similar cancer types show higher cosine similarities.

To quantitatively assess this clustering, we also found that *BulkRNABert* ‘s embedding space allows us to cluster similar cancer types more accurately. More precisely, we computed the cosine similarity between every couple of raw RNA-seq samples of each cohort and compare them to the cosine similarity between every couple of RNA-seq embeddings of each cohort and got a relative increase of 36% in cosine similarity when using embeddings.

### 3.5. Survival analysis

The performance analysis of *BulkRNABert* on survival prediction is divided into two parts: first, we consider pan-cancer models, and then we dig into 5 per-cohort survival analyses.

#### 3.5.1. Pan-cancer analysis

##### Experimental setup

For the pan-cancer analysis, we trained survival models (same architecture as the classification head, with a slightly larger MLP with hidden layers sizes of [512, 256]), based on the different pre-trained language models. The pan-cancer dataset is composed of 11,035 samples, split into train/test following the same experimental methodology as for the classification task.

##### Results

We provide in Table 2 the pan-cancer survival results, comparing with *CustOmics* (Benkirane et al., 2023) and with Masking Auto-Encoder (MAE) from Gross et al. (2024) inspired by a method designed to learn representations of tabular data (Yoon et al., 2020). Our models are evaluated using Harrell’s C-index (Harrell et al., 1982). For the pan-cancer setting, we report two different C-indexes: C-index on the whole test set (all cohorts) referred to as “C-index”, and a weighted sum of the C-indexes computed per cohort on the pan-cancer test set (with weights corresponding to the number of samples of each cohort in the test set), referred to as “Weighted C-index”. The motivation behind providing these 2 metrics was to confirm that the model was not only learning to differentiate between cancer types without being able to correctly predict survival time within a given cohort. We also provide in Appendix C (Table 7) the non-weighted average C-index (“Macro C-index”). Overall, our method exhibits state-of-theart results using the language model pre-trained on TCGA. One notices that the survival model built on top of *BulkRNABert(GTEx ENCODE)* provides results close to the baselines we are comparing with, thus highlighting the effectiveness of the pre-training. We keep in mind that *CustOmics* reaches a weighted test C-index of 0.68 when adding other omics sources, paving the way for further improvement.

**Table 2:** Pan-cancer survival analysis.

| Methods | C-index | Weighted C-index |
| --- | --- | --- |
| *CustOmics* (RNA-seq only) Benkirane et al. (2023) | NA | $0.630 \pm 0.020$ |
| MAE Gross et al. (2024) | $0.756 \pm 0.010$ | NA |
| *BulkRNABert*(GTEx ENCODE) | $0.753 \pm 0.009$ | $0.621 \pm 0.015$ |
| *BulkRNABert*(TCGA) | **$0.765 \pm 0.011$** | **$0.642 \pm 0.014$** |

#### 3.5.2. Per-cohort analysis

##### Experimental setup

The same analysis is performed by considering survival models trained on specific cohorts: GBMLGG, LUAD, UCEC, BLCA, BRCA. Due to the smaller number of samples available per cohort compared to the pan-cancer setting, training a model for a particular cohort is a harder problem (see Appendix C for the exact number of samples per cohort in the survival setting).

##### Results

We compare in Table 3 our method to the same baselines (*CustOmic* and MAE), to which we add a comparison with a simple MLP (same architecture as the one used on top of the language model) trained on raw RNA-seq data, which proves to be the best on BRCA. However, our method (using pre-training on TCGA) reaches state-of-the-art results for the BLCA and LUAD cohorts, and brings a significant improvement on GBMLGG and UCEC cohorts. Our model even reaches better results than *CustOmics* trained on multiple omics sources on 3 cohorts.

**Table 3:** Per-cohort survival analysis.

| Methods | GBMLGG | LUAD | UCEC | BLCA | BRCA |
| --- | --- | --- | --- | --- | --- |
| Baseline | $0.831 \pm 0.016$ | $0.550 \pm 0.053$ | $0.609 \pm 0.065$ | $0.598 \pm 0.034$ | **$0.676 \pm 0.057$** |
| *CustOmics (multi-omics)* | $0.642 \pm 0.028$ | $0.625 \pm 0.037$ | $0.667 \pm 0.022$ | **$0.637 \pm 0.050$** | $0.633 \pm 0.018$ |
| *CustOmics (RNA-seq only)* | $0.609 \pm 0.057$ | $0.638 \pm 0.040$ | $0.660 \pm 0.032$ | $0.626 \pm 0.071$ | $0.611 \pm 0.022$ |
| MAE | $0.775 \pm 0.038$ | $0.621 \pm 0.070$ | $0.633 \pm 0.071$ | $0.615 \pm 0.039$ | $0.632 \pm 0.044$ |
| *BulkRNABert(TCGA)* | **$0.844 \pm 0.023$** | **$0.648 \pm 0.057$** | **$0.703 \pm 0.040$** | $0.627 \pm 0.069$ | $0.604 \pm 0.032$ |

**Table 4:** Influence of pre-training.

| Model | test macro-F1 |
| --- | --- |
| Pre-trained (TCGA) | **0.918 ± 0.006** |
| Pre-trained (GTEx ENCODE) | 0.914 ± 0.007 |
| No pre-training + full fine-tuning | 0.894 ± 0.014 |
| Pre-training + full fine-tuning | 0.909 ± 0.008 |

#### 3.5.3. Transfer learning from pan-cancer to per-cohort

##### Experimental setup

We conducted a final experiment to further attest the performance of pan-cancer models on particular cohorts. To this end, we compare 3 models for a given cohort as shown in Figure 5: a per-cohort survival model trained on that particular cohort only from a language model pre-trained on TCGA, a pan-cancer model (*BulkRNABert(TCGA)*) from which we only report the C-index on its test set restricted to the cohort of interest, and finally the same pan-cancer model but whose training set has been pruned of any samples of the cohort of interest. This experiment has been carried out on the 5 previous cohorts and on the 4 smallest TCGA cohorts.

**Figure 5:**
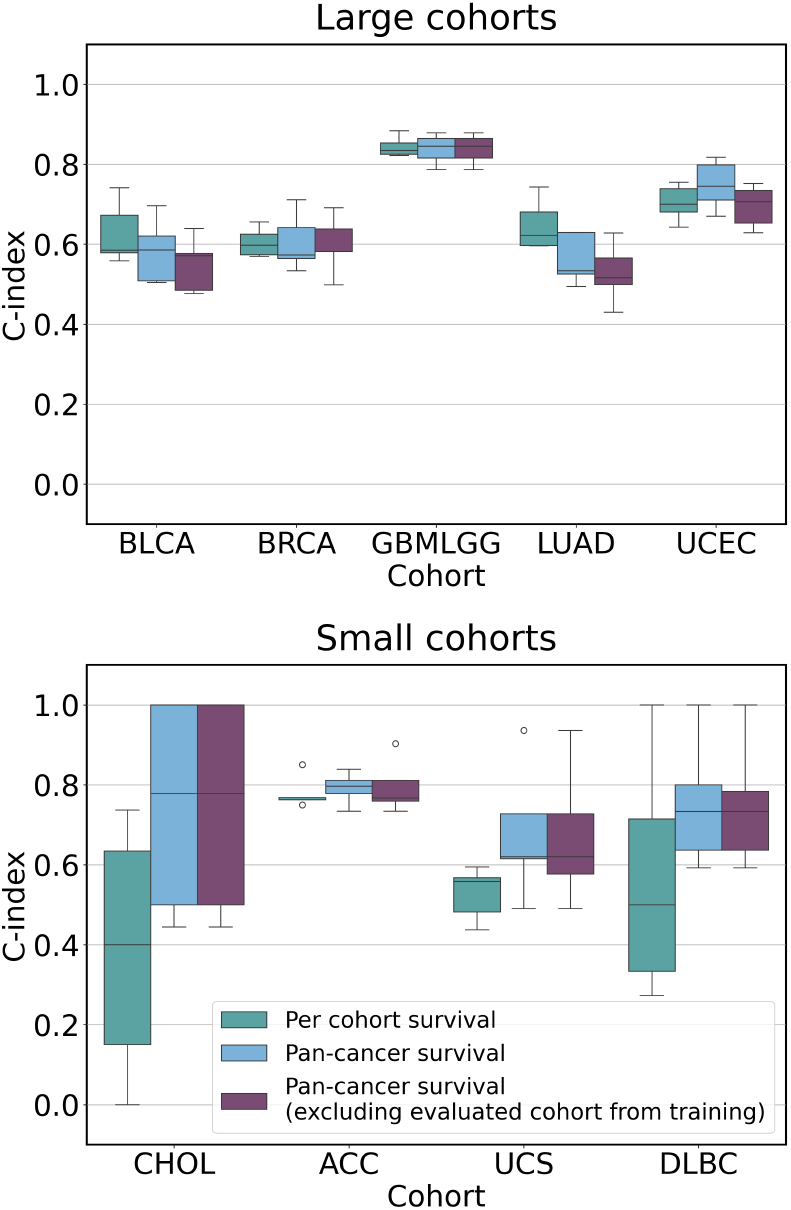
Comparison of per-cohort and pan-cancer survival models evaluated on selected cohorts. 1st box plot: per-cohort model. 2nd box plot: pan-cancer model (BulkRN-ABert(TCGA). 3rd box plot: pan-cancer model with evaluated cohort excluded from the training set.

##### Results

For the 5 first cohorts, pan-cancer models, though sometimes providing slightly worse scores, reach C-indexes that are very close to the per-cohort setting, even when the cohort of interest had been removed from the training set: this shows transfer learning capabilities of our method by being able to infer survival times on unseen cohorts. Finally, for the UCEC cohort, an improvement is observable from the per-cohort to the pan-cancer model, which may be rooted in the fact that UCEC is a highly censored dataset (only 16% of non-censored data). The same conclusion applies to the 4 smallest cohorts, for which one makes a significant improvement when considering pan-cancer models, showing that survival signals from other cohorts may benefit others.

## 4. Conclusion

In this work, we present *BulkRNABert*, the first language model pre-trained on bulk RNA-seq data and fine-tuned on cancer type classification and survival analysis. We came up with new state-of-the-art results on both tasks, with significant improvements in the pan-cancer settings, showing the strength of our pre-training method. We additionally showcase the transfer learning capabilities of our pan-cancer survival models to unseen cohorts. Further improvements may involve refining the results in the per-cohort setting, especially trying to understand the grounding of the great performance of baselines for the BRCA cohort. This may call for leveraging some interpretation methods, in particular quantifying the importance of each gene in survival predictions. Also, as shown in Benkirane et al. (2023), the addition of other omics sources exhibits an increase in performance on both downstream tasks. We are thus interested in leveraging genomics and imaging data to consider multi-modal models. Finally, though only pretraining with non-cancerous RNA-seq data already provides models that go head to head with state-of-the-art methods, we did not observe further improvement by mixing these data with cancerous data compared to models directly pre-trained on TCGA. Scaling up the language models could help cope with it.

## Acknowledgments

Research supported by Cloud TPUs from Google’s TPU Research Cloud (TRC).

## Appendix A. Language model pre-training

In Figure 6, we compare the true distributions of tokens on a few samples with the predicted distributions by *BulkRNABert*. One can see that the model correctly learns RNA-seq token distribution.

**Figure 6:**
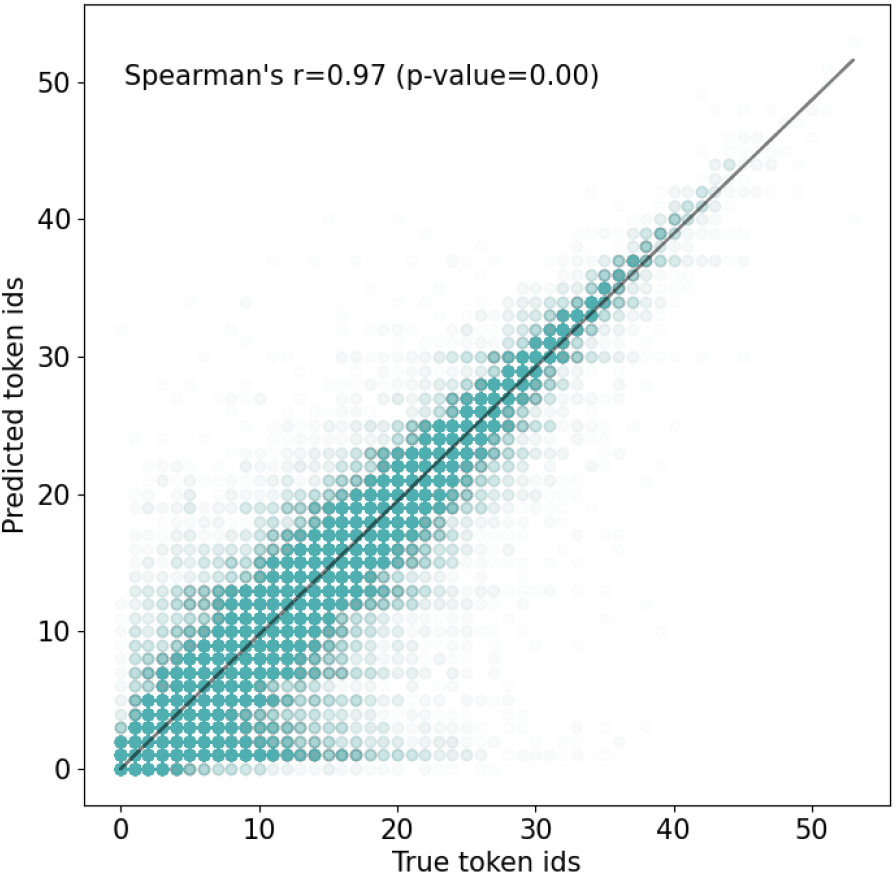
True vs. predicted tokens distribution using *BulkRNABert(TCGA)*.

In Figure 7, we provide the learning curves, in terms of reconstruction accuracy, of the masked language modeling task for the 3 *BulkRN-ABert* models corresponding to the 3 pre-training datasets: *BulkRNABert(GTEx ENCODE), BulkRN-ABert(GTEx ENCODE TCGA)*, and *BulkRN-ABert(TCGA)*.

**Figure 7:**
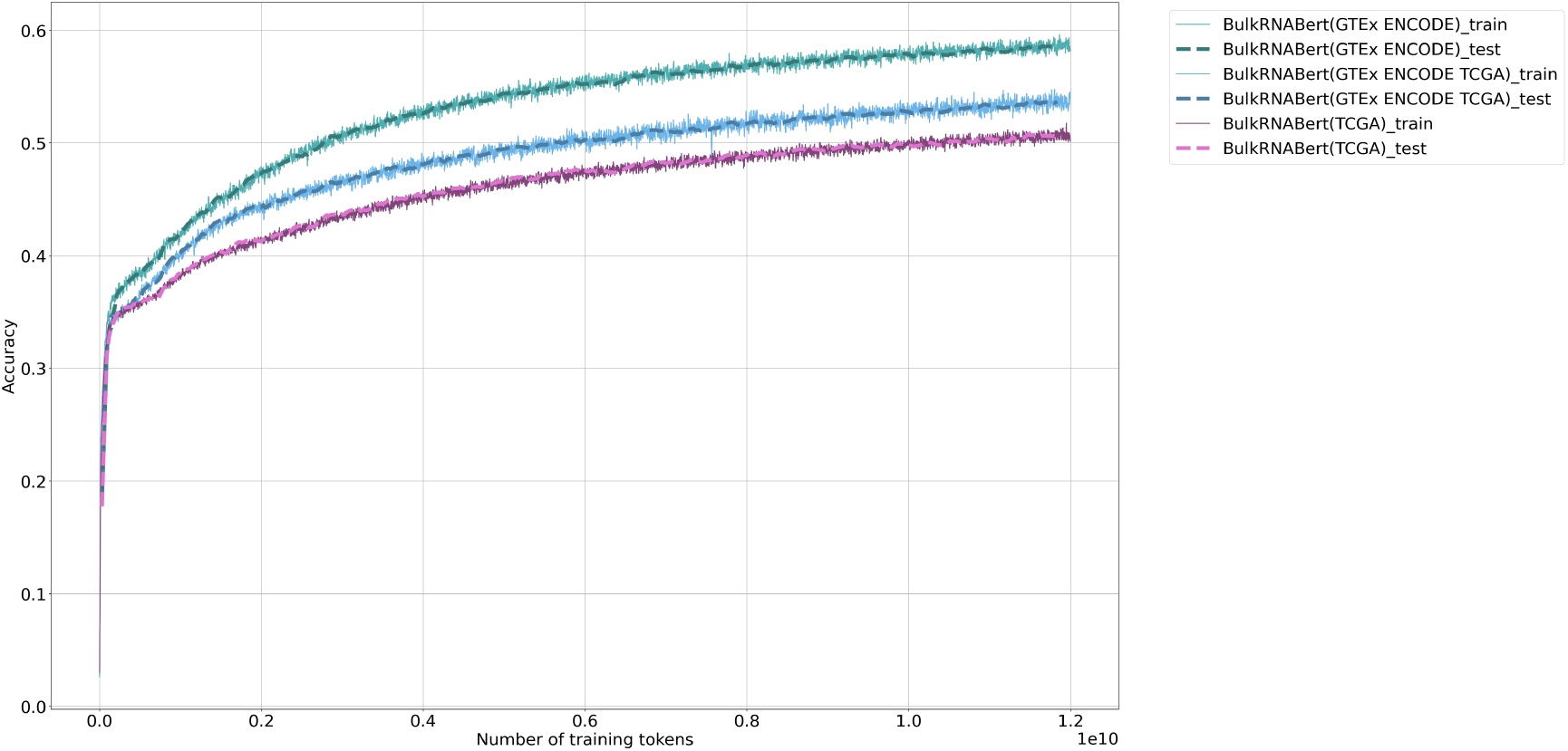
BulkRNABert pre-training with Masked Language Modeling on 3 different datasets. One observes higher reconstruction accuracy on non-cancerous tissues (GTEx ENCODE) than cancerous tissues (TCGA).

## Appendix B. Cancer type classification

For the cancer type classification task, we report the label distribution for both the 5 cohorts setting (Figure 8) and the pan-cancer setting (Figure 9). The latter shows a high class imbalance, hence the need to report both weighted and macro *F*_1_ scores. We furthermore report in Table 5 the performance of *BulkRNABert* on the 5 cohorts classification problem, completed with the results where we only train an MLP without using the *IA*^3^ method. We thus observe a slight improvement in mean weighted *F*_1_ score when using the parameter-efficient method. An ablation study on the impact of the dataset used to pre-train *BulkRNABert* is also provided in Table 6 for the pan-cancer classification task. Finally, a confusion matrix for the pan-cancer setting is provided in Figure 10 for the *BulkRNABert(TCGA) + MLP + IA*^3^ model, showing great classification performance with errors mainly between very similar cancer types (*e*.*g* between COAD - colon cancer - and READ - rectum cancer). An ablation study on the influence of pre-training is provided in 4 in which we compared pan-cancer classification models built on top of pretrained language models (either on TCGA or GTEx and ENCODE) or a randomly initialized language model which is fully fine-tuned (cf 3.3 for the details of the models).

**Table 5:** 5 cohorts classification complete results. “*BulkRNABert(Pre-Training Dataset)* + MLP/SVM” means that the MLP/SVM is trained, while freezing the language model weights. Adding “+ *IA*^3^” indicates that the language models’ weights are fine-tuned using the *IA*^3^ method. *m* for PCA and NMF indicates the number of components.

| Methods | test weighted-F1 |
| --- | --- |
| NMF(m=128) + SVM | 0.898 ± 0.018 |
| PCA(m=256) + SVM | 0.968 ± 0.010 |
| *CustOmics (RNA-seq only)* Benkirane et al. (2023) | 0.946 ± 0.011 |
| *CustOmics (multi-omics)* Benkirane et al. (2023) | 0.964 ± 0.004 |
| *BulkRNABert(GTEX ENCODE)* + SVM | 0.977 ± 0.008 |
| *BulkRNABert(GTEX ENCODE)* + MLP | 0.965 ± 0.010 |
| *BulkRNABert(GTEX ENCODE)* + MLP + $IA^3$ | 0.982 ± 0.008 |
| *BulkRNABert(GTEX ENCODE TCGA)* + SVM | 0.986 ± 0.004 |
| *BulkRNABert(GTEX ENCODE TCGA)* + MLP | 0.982 ± 0.006 |
| *BulkRNABert(GTEX ENCODE TCGA)* + MLP + $IA^3$ | 0.989 ± 0.004 |
| *BulkRNABert(TCGA)* + SVM | 0.991 ± 0.005 |
| *BulkRNABert(TCGA)* + MLP | 0.991 ± 0.004 |
| *BulkRNABert(TCGA)* + MLP + $IA^3$ | **0.993 ± 0.004** |

**Table 6:**
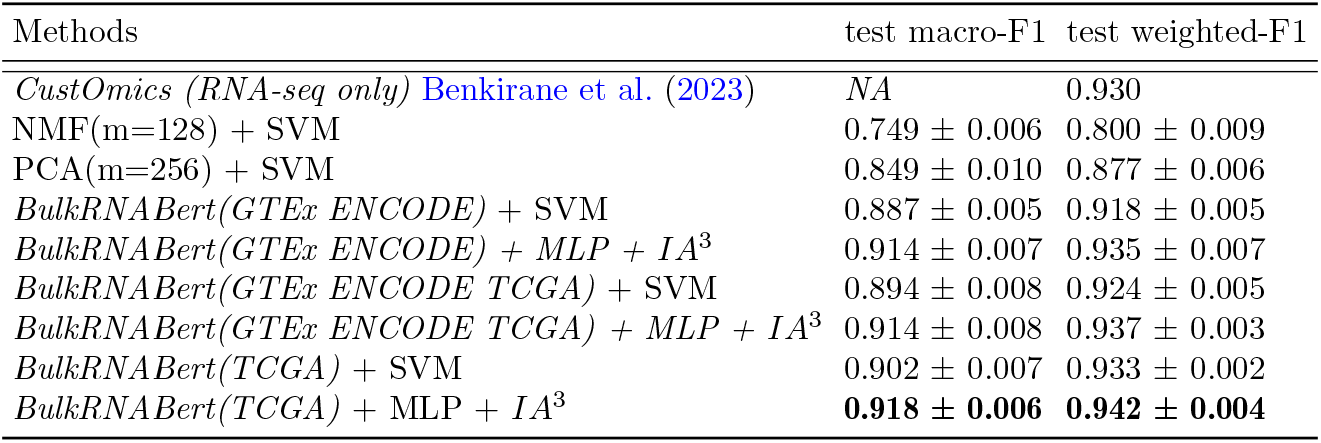
Pan-cancer classification. *NA* indicates that the score is not provided by the author.

**Figure 8:**
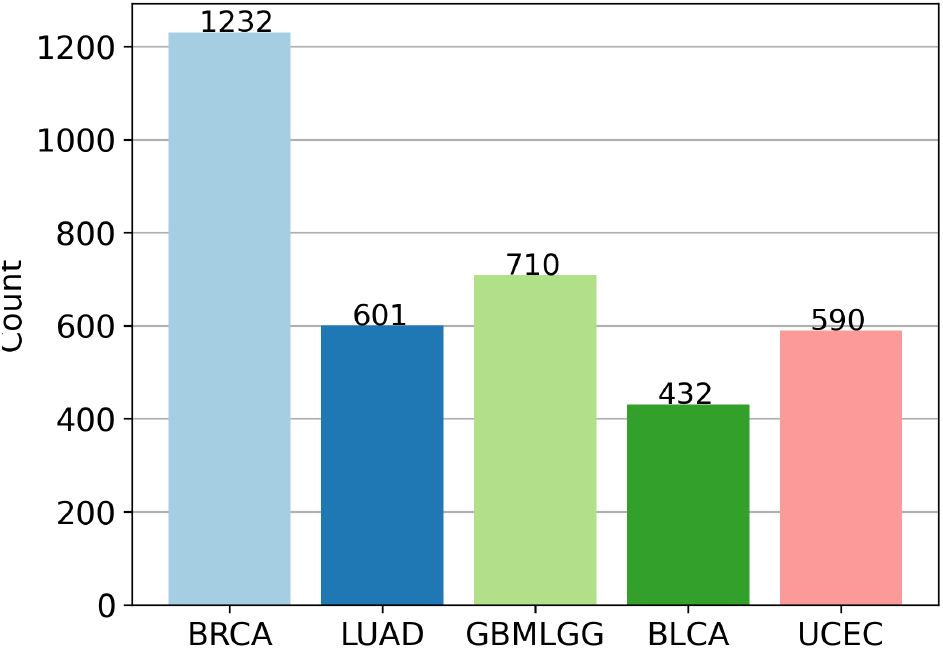
5 cohorts classification label distribution.

## Appendix C. Survival Analysis

We additionally provide in Figure 11 the number of available samples in the TCGA datasets for survival analysis for the specific cohorts we are interested in, both for the 5 large cohorts that guided our study (BRCA, BLCA, GBMLGG, LUAD, and UCEC), and for the 4 smallest TCGA cohorts (CHOL, ACC, UCS, and DLBC). Finally, we present again in Table 7 the performance of the pan-cancer survival models, adding the “Macro C-index” corresponding to the mean of the C-indexes computed by cohort on the test set of the pan-cancer dataset.

**Figure 9:**
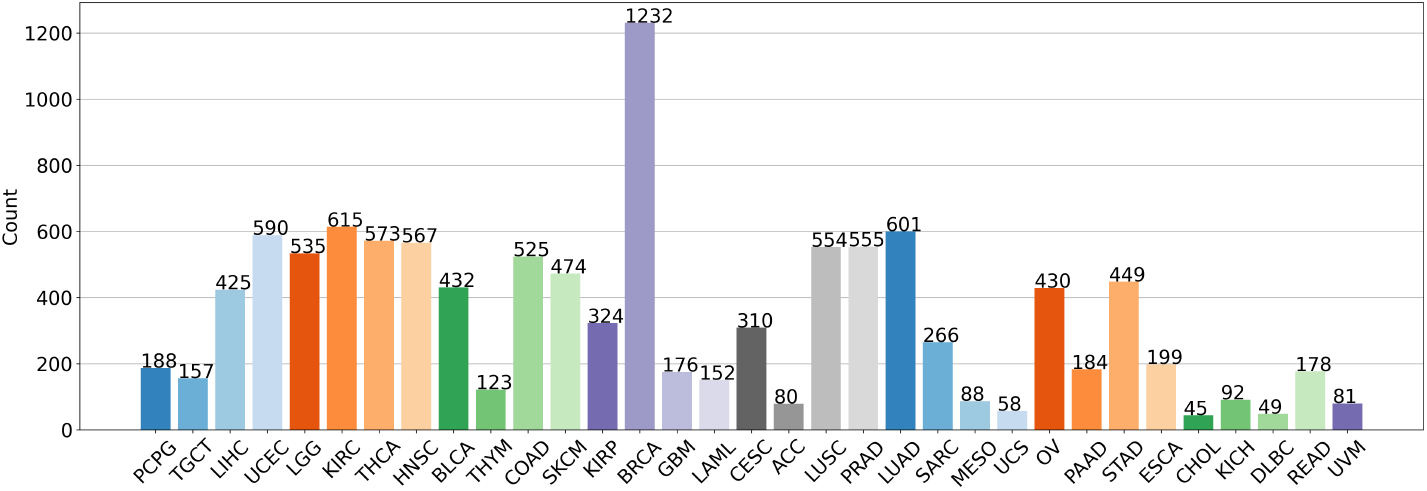
Pan-pancancer classification label distribution.

**Figure 10:**
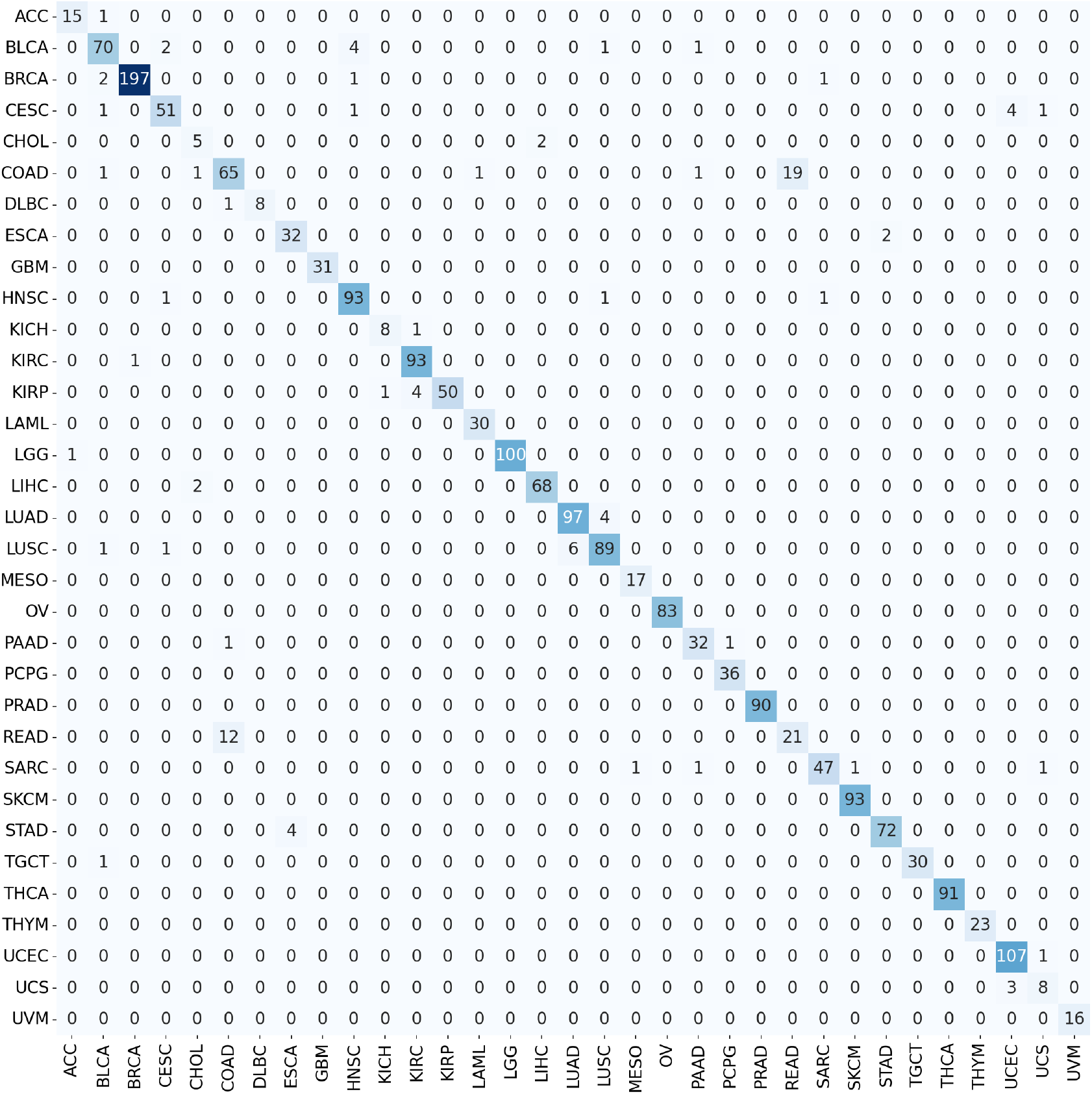
Confusion matrix for the pan-cancer classification problem for the model *BulkRNABert(TCGA) + MLP + IA*^3^.

**Figure 11:**
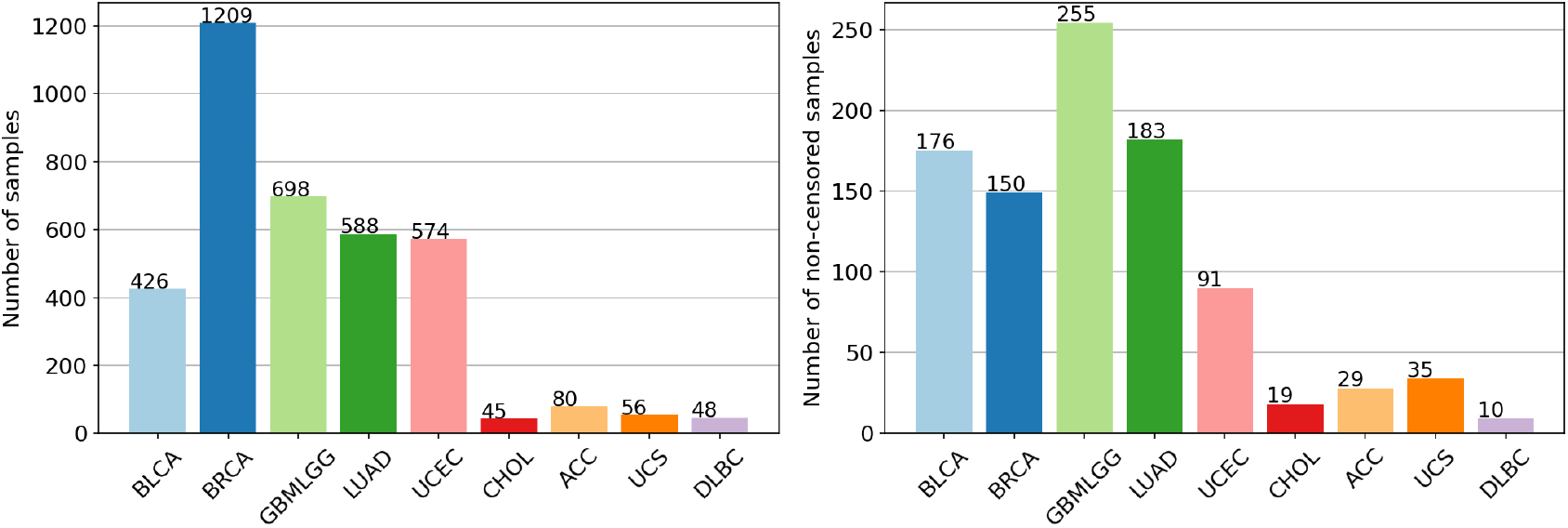
Per-cohort survival analysis: number of samples per cohort (left), and number of non-censored samples per cohort (right).

**Table 7:**
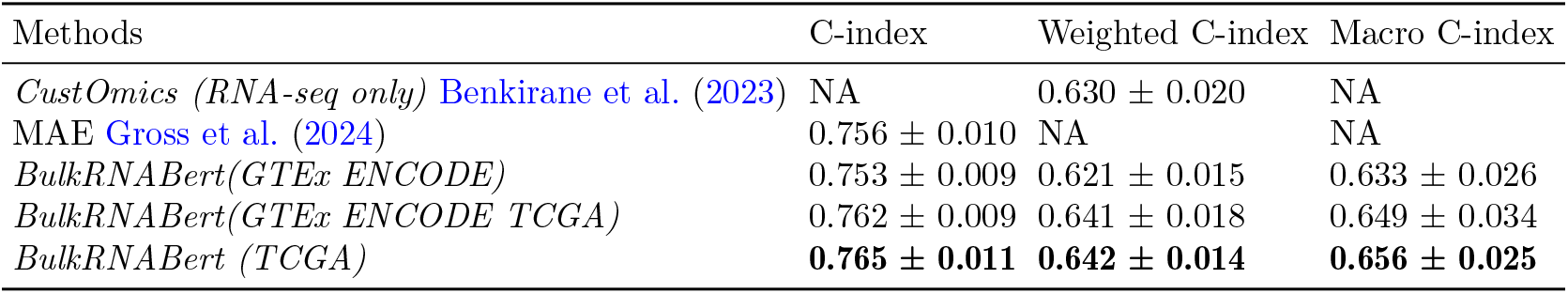
Pan-cancer survival analysis.

## Appendix D. Robustness to missing gene expressions

A common clinical issue when it comes to applying models like *BulkRNABert* which are pre-trained on a fixed set of genes, is that not all gene expressions might be available. To this end, we performed a study to understand the influence of missing gene expressions. To do so, on our pan-cancer classification dataset, we randomly removed a certain percentage of gene expression from all samples and filled in the missing values with *BulkRNABert* predictions. We then analyzed the drop in performance of our best pan-cancer classification model as a function of the percentage of missing gene expressions. In Figure 12 is reported this performance ratio, showing that by imputing missing gene expressions using *BulkRN-ABert*, our classification model keeps most of its performance (the loss in performance is always less than the corruption rate), thus it is robust to missing data.

**Figure 12:**
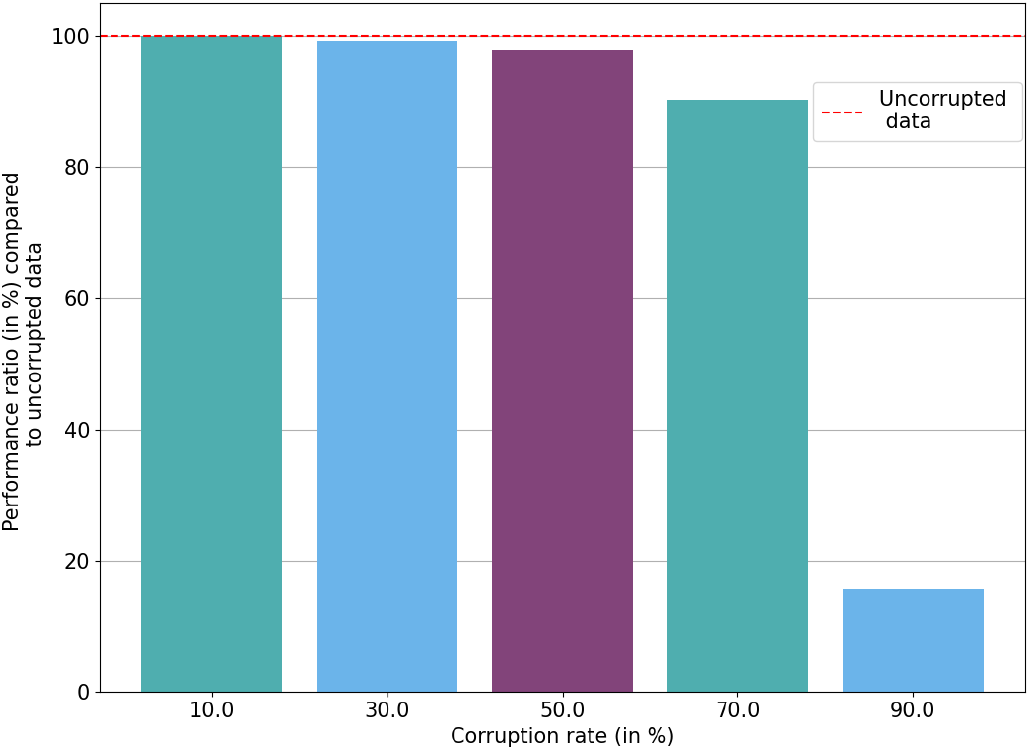
Performance ratio (in test macro F1-score) of pan-cancer classification model between original dataset and corrupted data (missing values inferred by *BulkRNABert*)

## Appendix E. TCGA cohorts abbreviations

We provide in Table 8 the abbreviations of the TCGA cohorts used throughout this paper.

**Table 8:** TCGA cohort abbreviations.

| Abbreviation | Name |
| --- | --- |
| BLCA | Bladder Urothelial Carcinoma |
| BRCA | Breast invasive carcinoma |
| GBM | Glioblastoma multiforme |
| LGG | Brain Lower Grade Glioma |
| LUAD | Lung adenocarcinoma |
| UCEC | Uterine Corpus Endometrial Carcinoma |
| CHOL | Cholangiocarcinoma |
| ACC | Adrenocortical carcinoma |
| UCS | Uterine Carcinosarcoma |
| DLBC | Lymphoid Neoplasm Diffuse Large B-cell Lymphoma |

## Notes

### Competing Interest Statement

centralesupelec.fr, instadeep.com

### Summary of Updates

Update after submission to ML4H 2024.

